# A Query-to-Dashboard Framework for Reproducible PubMed-Scale Bibliometrics and Trend Intelligence

**DOI:** 10.64898/2026.02.27.708328

**Authors:** Benjamin L. Kidder

**Author notes:** Correspondence: Benjamin L. Kidder.

## Abstract

The rapid expansion of biomedical literature necessitates computational approaches for systematic analysis of publication patterns, identification of emerging scientific themes, and characterization of field evolution. We present PubMed Atlas, an integrated command-line and web-based platform for conducting topic-specific bibliometric analyses through programmatic access to PubMed E-utilities. This workflow retrieves PubMed identifiers matching user-defined queries, downloads comprehensive metadata in batch mode, extracts structured information including titles, abstracts, author affiliations, Medical Subject Headings, publication classifications, funding acknowledgments, and digital object identifiers, then organizes these data within a local SQLite relational database optimized for rapid queries and visualization. An accompanying Streamlit-based interactive dashboard enables exploration of temporal publication patterns, journal distribution profiles, MeSH term frequencies, geographic author distributions, and direct linking to recent publications. We demonstrate the application of PubMed Atlas to cancer stem cell biology and stem cell transcriptional regulatory network research, providing a framework for reproducible bibliometric investigation and systematic identification of research gaps within dynamically evolving scientific domains.

## INTRODUCTION

PubMed’s biomedical literature index has experienced substantial expansion throughout the past twenty years, encompassing more than 36 million citations by 2024^1^. This remarkable growth creates simultaneous opportunities and obstacles for investigators studying stem cell biology and cancer: although unprecedented volumes of information have become accessible, the tasks of locating pertinent publications, monitoring evolving trends, and comprehending research landscapes have grown substantially more complex^2,3^. Conventional approaches to literature synthesis, despite their continued utility, demand intensive manual effort, may reflect biases inherent to reviewer backgrounds, and present difficulties for incorporating newly published findings^4,5^.

Quantitative bibliometric methodologies offer complementary strategies to traditional narrative reviews through systematic examination of publication dynamics, citation relationships, collaborative author networks, and temporal research patterns across extensive document collections^6,7^. These analytical frameworks can illuminate the organizational structure and temporal evolution of scientific disciplines, distinguish particularly influential contributions and investigators, characterize collaborative networks, reveal nascent research directions, and inform strategic planning for research initiatives and funding allocation^8^. Within stem cell research, bibliometric investigations have chronicled the rapid proliferation of induced pluripotent stem cell (iPSC) research subsequent to the groundbreaking contributions of Takahashi and Yamanaka in 2006^9^, documented the developmental trajectory of organoid methodologies^10^, and examined publication dynamics within cancer stem cell research^11,12^.

Notwithstanding the demonstrated utility of bibliometric investigation, most biological researchers have limited access to specialized analytical platforms or lack computational proficiency necessary for developing customized analytical solutions. Commercial database platforms including Web of Science, Scopus, and Dimensions deliver bibliometric functionality but necessitate institutional licensing agreements and may lack adequate flexibility for discipline-specific investigations^13,14^. Open-source visualization platforms such as VOSviewer and Bibliometrix deliver robust analytical capabilities but require independent data procurement workflows^15,16^. Additionally, numerous existing platforms fail to provide cohesive integration across data acquisition, storage, computational analysis, and visualization components.

PubMed, curated by the National Library of Medicine (NLM), constitutes the principal repository for biomedical literature and delivers unrestricted access through the Entrez information retrieval system^17^. The E-utilities (Entrez Programming Utilities) Application Programming Interface (API) facilitates programmatic data access, enabling investigators to execute searches, obtain metadata, and retrieve comprehensive records^18^. However, effective utilization of E-utilities necessitates comprehension of RESTful API architecture, XML document parsing, rate constraint policies, and database schema design, technical competencies that constitute substantial barriers for numerous biological investigators.

PubMed Atlas addresses these technical obstacles by delivering an integrated, open-source analytical workflow connecting E-utilities data procurement with local database architecture and interactive data visualization. The platform constructs a persistent “literature atlas” permitting repeated interrogation without redundant API requests, accommodates user-defined topic specifications such as stem cell and cancer research, incorporates automated metadata extraction with normalization, executes geographic inference from institutional affiliations, computes bibliometric indicators including compound growth rates and temporal moving averages, and delivers an intuitive web-based interface requiring no programming expertise. Unlike visualization-only platforms (e.g., VOSviewer^15^) or analysis libraries that require manual data preprocessing (e.g., Bibliometrix^7^), PubMed Atlas integrates end-to-end acquisition, normalization, persistent storage, reproducible configuration, and interactive exploration within a single local, version-controlled framework.

The architectural design of PubMed Atlas embodies established best practices for scientific software engineering, incorporating modular design with functional separation of API communication, document parsing, database operations, and analytical computation; comprehensive error management and retry mechanisms for robust API interactions; optional response caching to minimize redundant network requests; rate constraint compliance conforming to NCBI policies; and reproducible analytical workflows through structured configuration files (**Figure 1**). The SQLite database schema implements normalized relational tables supporting efficient query execution while preserving data integrity.

**Figure 1.**
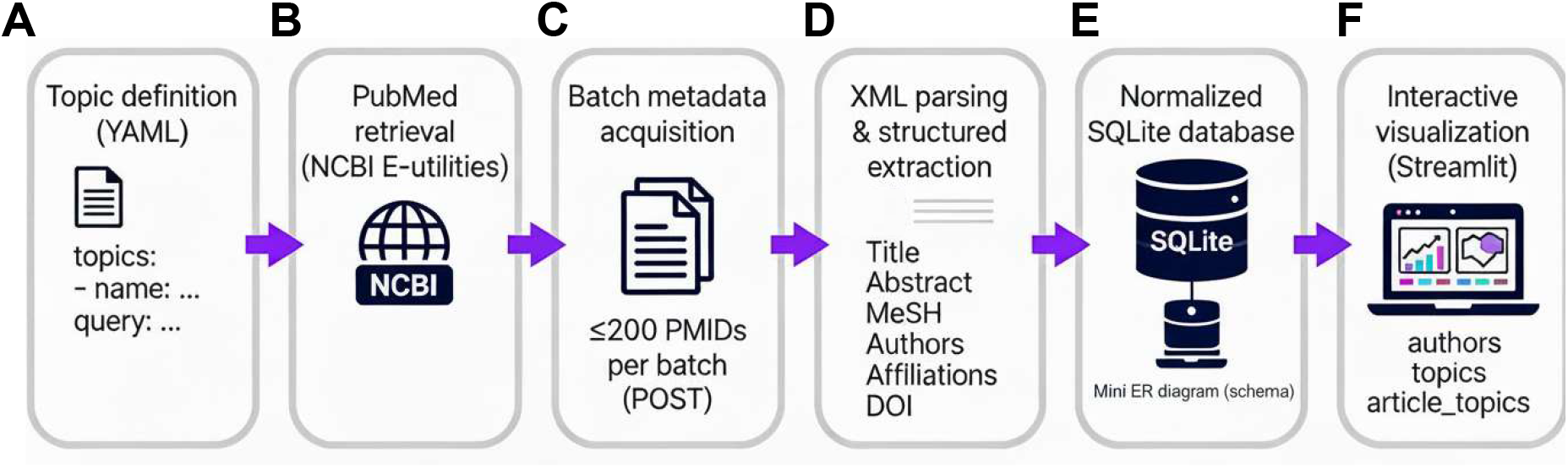
PubMed Atlas Workflow Architecture. Schematic diagram illustrating the modular architecture of PubMed Atlas. The workflow encompasses (A) topic specification through YAML configuration files, (B) programmatic PubMed access via NCBI E-utilities API, (C) batch metadata acquisition through EFetch requests, (D) XML parsing and structured data extraction via lxml, (E) normalized storage within SQLite relational database, and (F) interactive visualization through Streamlit-based web dashboard. Error handling mechanisms, rate limiting compliance, and optional response caching ensure robust and reproducible operations.

This comprehensive protocol details the complete operational workflow for utilizing PubMed Atlas to execute bibliometric investigations relevant to topics such as stem cell transcriptional networks and cancer stem cell biology. We present procedures for software installation, configuration of research-specific topics, execution of searches to construct literature databases, analysis of publication trends and quantitative metrics, exploration through the interactive visualization dashboard, and exportation of results for publication.

The framework described here is applicable to any research domain indexed within PubMed and can be adapted to address specific bibliometric research questions and trend intelligence requirements.

## RESULTS

### Validation using representative biomedical domains

We evaluated PubMed Atlas using representative biomedical research domains spanning stem cell transcriptional networks, cancer stem cell biology, and cellular reprogramming. Queries were defined using PubMed-compatible Boolean syntax and executed through programmatic access to the Entrez E-utilities API. For the benchmarked cancer query (**Table S1**), total wall time for retrieval and ingestion of 500 records was 11.56 seconds. The majority of execution time is attributable to batched EFetch metadata retrieval (200 records per POST request), while identifier retrieval via ESearch contributed minimally to overall runtime. No retrieval failures were observed during benchmark execution.

Performance scaled predictably with record count under authenticated API access, with execution time reflecting expected external rate constraints. Execution time reflected expected external rate constraints rather than local computational overhead, indicating stable and reproducible behavior across moderate-scale retrieval tasks.

### Database storage characteristics and query performance

Normalized metadata were persisted within a relational SQLite database, enabling efficient aggregation and cross-topic reuse. Database size scaled linearly with article count, stabilizing at approximately 4.55–4.63 kilobytes per article across benchmark datasets ranging from 500 to 5,000 records (Table S1). For example, a database containing 5,000 records occupied 22.62 megabytes of disk storage.

Indexed aggregation queries executed with low latency on benchmark datasets. Persistent storage eliminated redundant API calls during iterative analyses. Re-execution of metric computation or cross-topic comparisons on previously retrieved datasets completed in under 2 seconds, representing a substantial efficiency gain relative to repeated API-based retrieval.

### Bibliometric metric computation

Analytical routines computed descriptive bibliometric metrics directly from the normalized database. For the pluripotency network dataset (n = 450), annual publication counts demonstrated steady growth across the past decade, with a calculated compound annual growth rate (CAGR) of 8.34 percent and a mean annual growth rate of 12.5 percent. Temporal smoothing using a three-year moving average reduced short-term variability without altering long-term growth trends.

Journal frequency analysis identified Cell Stem Cell (n = 45), Nature (n = 38), Cell (n = 32), and Stem Cells (n = 28) as predominant publication venues within the selected dataset. MeSH descriptor aggregation revealed Pluripotent Stem Cells (n = 425), Transcription Factors (n = 312), and Gene Regulatory Networks (n = 287) as the most frequently annotated conceptual themes. Complete metric computation for datasets in the thousands of records completed in seconds on a standard workstation (Intel i7 CPU, 16 GB RAM).

Geographic assignment derived from affiliation parsing indicated United States predominance within pluripotency network publications, followed by China, the United Kingdom, Japan, and Germany. Geographic aggregation executed in seconds on benchmark datasets, demonstrating minimal computational overhead relative to database size.

### Cross-topic comparative analysis

To evaluate comparative analytical capability, identical metric pipelines were applied to multiple research topics. Cancer stem cell organoid research exhibited a higher calculated CAGR (15.2 percent) relative to chromatin-focused stem cell investigations (6.8 percent) over comparable time windows. Notably, organoid-focused cancer stem cell research (**Fig. 2**) exhibited a marked post-2016 acceleration relative to chromatin-focused investigations. Journal distribution profiles differed across topics, with translational oncology journals more prominently represented in organoid-focused datasets. Geographic distributions also varied, with broader international representation observed in rapidly expanding subdomains.

**Figure 2.**
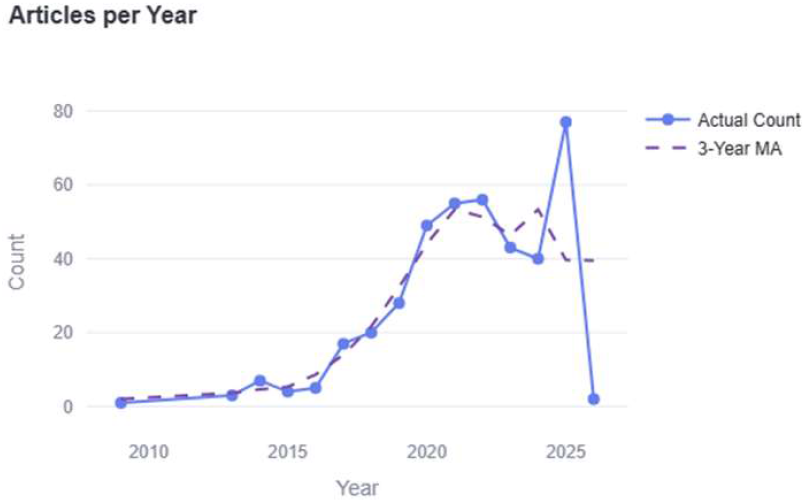
Temporal Publication Trends. Representative visualization of annual publication dynamics generated through PubMed Atlas analysis module. The line plot displays raw annual article counts (solid line) and three-year moving average (dashed line) for a selected research topic. Moving average calculation smooths short-term fluctuations while highlighting sustained growth patterns. Data derive from publication year metadata extracted from PubMed records and stored within SQLite database.

### Interactive visualization performance

The interactive dashboard rendered publication trends, journal distributions, MeSH frequencies, geographic summaries, and recent article tables using Plotly within a Streamlit interface (**Fig. 2-3**). For datasets up to 10,000 records, figure rendering latency remained below 1 second per visualization panel. Topic switching triggered dynamic re-aggregation of metrics with minimal perceptible delay, supported by indexed database queries.

**Figure 3.**
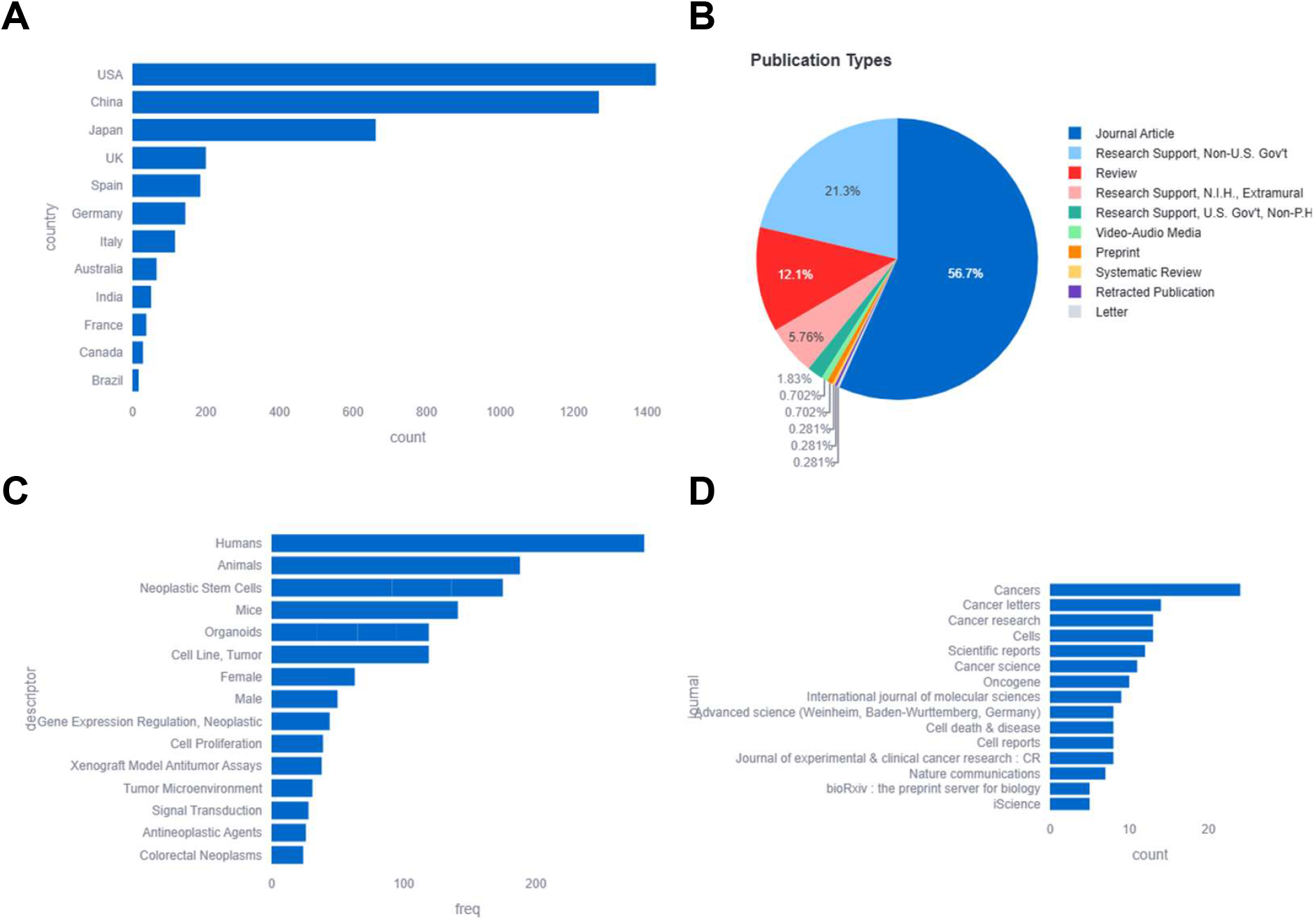
Interactive Dashboard Multi-Panel Visualization. Representative Streamlit dashboard output illustrating integrated bibliometric analyses. (A) Geographic distribution: top countries ranked by publication frequency inferred from author institutional affiliations with confidence-weighted assignment. (B) Publication type distribution: relative proportions of PubMed document classifications (Journal Article, Review, Clinical Trial, etc.). (C) MeSH term frequency: top Medical Subject Heading descriptors ranked by annotation frequency. (D) Journal distribution: top publication venues ranked by article counts. All panels dynamically update upon topic selection and support data export functionality.

Choropleth geographic visualizations scaled with country-level aggregation rather than raw article count, maintaining consistent responsiveness. Export functionality enabled figure download and CSV extraction of aggregated tables without requiring re-querying of external data sources.

### Reproducibility characteristics

Topic-specific datasets can be exported through direct SQLite queries to CSV format for external statistical analysis. To ensure reproducibility, analyses should document the PubMed Atlas version, complete query strings, date of data acquisition, maximum result limits, applied date filters, and database file checksum (MD5 or SHA256). Data manipulation and aggregation are implemented using pandas, relational queries utilize sqlite3, and visualization rendering uses Plotly. Metric computation scales approximately linearly with the number of retrieved records, with database indexing supporting efficient query execution for moderate-scale bibliometric datasets.

### Retrieval and ingestion benchmark performance

Using the PubMed query cancer AND medline[sb] AND hasabstract[text] NOT books[filter] (3,814,288 total hits), we benchmarked end-to-end retrieval and SQLite database construction for 100, 500, 1,000, and 5,000 records (**Table S1**). Wall clock runtime increased from 4.86 seconds for 100 records to 11.56 seconds for 500 records and 19.58 seconds for 1,000 records, while the 5,000 record run completed in 19.50 seconds. Effective ingestion throughput increased from 20.58 records per second (100 records) to 256.41 records per second (5,000 records), reflecting batching efficiency and amortization of connection overhead at larger scales. Peak resident memory increased from 85.8 MB to 388.2 MB across these runs. SQLite database size increased proportionally from 0.53 MB (100 records) to 22.62 MB (5,000 records), corresponding to 4.63 KB per article at 5,000 records. Benchmarks were conducted on a standard PC workstation equipped with an Intel i7 CPU and 16 GB RAM.

### Comparison with existing bibliometric tools

PubMed Atlas was evaluated relative to the R package Bibliometrix, a widely used open-source framework for bibliometric analysis. Bibliometrix requires manual export of PubMed records followed by file import and dataframe conversion within an R environment, whereas PubMed Atlas performs direct E-utilities-based retrieval, automated normalization, and persistent storage in a relational SQLite database. PubMed Atlas enables deterministic re-analysis without repeated file parsing or re-import and supports topic-based configuration through version-controlled YAML files. While Bibliometrix provides advanced citation and co-citation network analyses when citation metadata are available, PubMed Atlas emphasizes reproducible query-driven literature infrastructure, persistent query reuse, and integrated interactive dashboard exploration without requiring R programming. Key architectural differences are summarized in **Table S2**.

## DISCUSSION

PubMed Atlas provides an integrated framework for biomedical bibliometric analysis by combining data acquisition, storage, computation, and visualization within a unified open-source platform. The demonstrated application to stem cell and cancer research domains illustrates the platform’s capacity to illuminate publication trends, identify research priorities, characterize geographic contributions, and reveal temporal evolution of scientific themes. These applications demonstrate the feasibility of programmatic E-utilities access for constructing comprehensive, queryable literature databases.

Beyond serving as a bibliometric utility, PubMed Atlas illustrates a generalizable architecture for reproducible literature intelligence systems. By coupling deterministic query specification with persistent relational storage and version-controlled configuration, the platform transforms PubMed from a transient search interface into a reproducible analytical substrate.

The architectural design of PubMed Atlas incorporates several advantages relative to existing bibliometric platforms. First, the persistent SQLite database architecture eliminates redundant API requests during iterative analyses, reducing network overhead and accelerating repeated interrogations. Second, the flexible topic-based configuration framework accommodates arbitrary research domains through straightforward YAML specification, enabling rapid deployment for novel bibliometric questions. Third, the integrated visualization dashboard delivers immediate interactive exploration without requiring external software dependencies or specialized data format conversions. Fourth, the normalized relational database schema supports complex analytical queries extending beyond the pre-defined dashboard visualizations, accommodating advanced investigations through direct SQL interrogation.

Application to stem cell transcriptional networks and cancer stem cell research illustrates the platform’s ability to characterize publication venues and thematic structure across distinct domains. The predominant publication venues identified through our analysis align with established high-impact journals recognized for stem cell and cancer research dissemination.

MeSH term frequency analysis provided insights into conceptual priorities within each research domain. These quantitative MeSH analyses complement qualitative literature review by providing objective characterization of research emphases and temporal theme evolution.

Several technical considerations merit discussion regarding PubMed Atlas implementation. Citation relationship analysis remains unavailable through E-utilities API, limiting network analysis to co-authorship patterns derived from shared publications. Author name disambiguation, a recognized challenge in bibliometric investigation, requires external resolution as the platform implements simple string matching for author identification. Geographic inference from institutional affiliations, while generally accurate for explicitly stated country information, may introduce uncertainties for institutions with ambiguous location descriptors or investigators with multiple concurrent affiliations.

Future development directions for PubMed Atlas include integration of citation network analysis through alternative data sources, implementation of author disambiguation algorithms incorporating ORCID identifiers and institutional affiliations, extension to additional literature databases beyond PubMed, incorporation of full-text analysis for open-access publications, and development of machine learning models for automated research trend prediction and emerging theme identification.

The reproducibility of bibliometric analyses conducted through PubMed Atlas merits emphasis. By preserving query specifications, temporal constraints, and maximum result limits within configuration files, and by documenting the specific software version employed, investigators can ensure independent verification of published bibliometric findings.

In summary, PubMed Atlas delivers an integrated, accessible platform for conducting rigorous bibliometric investigations of biomedical literature. The demonstrated applications to stem cell transcriptional networks and cancer stem cell research illustrate the platform’s capacity to illuminate research trends, characterize publication landscapes, and identify knowledge gaps within dynamically evolving scientific domains. The open-source distribution, comprehensive documentation, and flexible configuration framework position PubMed Atlas as a valuable resource for the biomedical research community, enabling quantitative literature analysis without requiring specialized programming expertise or commercial software licenses.

## MATERIALS & METHODS

### System architecture and implementation

PubMed Atlas is a Python-based framework for topic-driven bibliometric analysis of PubMed-indexed literature. The system follows a modular architecture consisting of query specification, programmatic data acquisition via NCBI E-utilities, XML parsing and normalization, relational database persistence, and analytical and visualization layers. The software is distributed through a public GitHub repository and managed using a Conda-based dependency environment. The command-line interface exposes three primary entry points: search for database construction, analyze for metric computation, and dashboard for interactive visualization. Core dependencies include requests for API communication, lxml for XML parsing, pandas for data manipulation, sqlite3 for database access, and Plotly and Streamlit for visualization rendering.

### Topic specification and query design

Research domains are defined through YAML configuration files that associate unique topic identifiers with PubMed-compatible query strings. Queries employ Boolean operators (AND, OR, NOT) and standard PubMed field tags including [Title/Abstract], [MeSH], [PDAT], [Author], and [Journal]. Temporal filtering is implemented using publication date constraints within the [PDAT] field. All queries are validated against the PubMed web interface prior to execution to confirm syntactic correctness and expected retrieval scope. Topic definitions are stored in the database and linked to retrieved articles through a join table to enable many-to-many mappings between articles and research domains.

### Data acquisition and normalization

Literature retrieval is performed using the Entrez E-utilities API. For each topic, an ESearch request retrieves PubMed identifiers (PMIDs) matching the specified query. Identifiers are subsequently submitted to EFetch in batches of up to 200 PMIDs per HTTP POST request to obtain complete XML metadata records. XML parsing is performed using lxml to extract structured bibliographic fields including PMID, title, abstract, journal, publication date, DOI, author names and affiliations, MeSH descriptors and qualifiers, publication types, grant agencies and identifiers, and author-supplied keywords. Extracted records are normalized and inserted into a SQLite relational database.

The database schema enforces referential integrity using primary and foreign key constraints. The articles table, keyed by PMID, stores core metadata. Related tables include authors, meshterms, publicationtypes, grants, keywords, and geography, each linked to articles by PMID. Topic definitions are stored in a topics table, and article-to-topic mappings are implemented through an articletopics join table. Primary key constraints prevent duplicate insertion of PMIDs while allowing individual articles to be associated with multiple topics without redundant storage of bibliographic metadata.

Geographic inference is performed using rule-based pattern matching on affiliation strings against curated country and city name dictionaries. Matching prioritizes explicit country mentions, followed by city-level inference when country names are absent. A heuristic confidence score between 0 and 1 is assigned based on match specificity. Geographic assignment is descriptive and not externally validated.

API usage complies with NCBI rate limits, maintaining a maximum of three requests per second for unauthenticated access and ten requests per second when an API key is supplied. Transient network failures or rate limit violations trigger retry logic with exponential backoff. Optional caching stores raw XML responses keyed by query parameters to reduce redundant API calls during repeated analyses.

### Bibliometric metric computation

Analytical routines operate directly on the normalized SQLite database using pandas-based aggregation. Computed metrics include total article counts per topic, annual publication counts, compound annual growth rate (CAGR), mean annual growth rate, journal frequency rankings, MeSH descriptor and descriptor-qualifier frequencies, and country-level publication counts. CAGR is calculated as (V_final / V_initial)^(1/n) − 1, where V_initial and V_final correspond to the first and last non-zero annual publication counts and n represents the number of intervening years. Years with zero counts are excluded from denominator calculations to avoid undefined growth estimates. Temporal smoothing is performed using a three-year moving average applied to annual counts. All analyses are descriptive and do not include inferential statistical testing.

### Visualization framework

Interactive visualization is implemented using Plotly within a Streamlit-based dashboard interface. The dashboard dynamically renders annual publication trends, journal distribution bar charts, MeSH term frequency distributions, choropleth world maps summarizing geographic patterns, and tabular views of recent articles. Visualizations update automatically based on selected topic identifiers and support export using Plotly’s native download functionality. Tabular data can be exported in CSV format for external analysis.

### Extended analytical capabilities

The relational database design supports structured downstream analyses. Author-level publication frequencies can be extracted using SQL queries to generate input files for external network visualization platforms such as Gephi or Cytoscape. Grant table aggregation enables ranking of funding agencies by frequency within a topic. Cross-topic comparisons are performed by applying identical growth and distribution metrics across multiple stored topics, enabling systematic evaluation of differences in temporal trends, journal representation, and geographic patterns without requiring re-fetching of metadata.

## Code availability

PubMed Atlas is implemented in Python and is freely available under an open-source license at: https://github.com/KidderLab/PubMed-Atlas. The repository includes source code, documentation, example configuration files, and environment specifications required to reproduce analyses.

## AUTHOR CONTRIBUTIONS

B.L.K. conceived of the study, performed data analysis, and drafted the manuscript.

## ACKNOWLEDGEMENTS

This work utilized the Wayne State University High Performance Computing Grid for computational resources (https://www.grid.wayne.edu/).

